# *XPO1* gene therapy restores cardiac function in rats with chronic induced myocardial infarction

**DOI:** 10.1101/575340

**Authors:** María García-Manzanares, Carolina Gil-Cayuela, Luis Martínez-Dolz, José Ramón González-Juanatey, Francisca Lago, Manuel Portolés, Esther Roselló-Lletí, Miguel Rivera

## Abstract

**Background:** In previous studies, we showed that several nuclear-cytoplasmic transport molecules were closely related to ventricular dysfunction in human heart failure. Particularly, the transcriptomic signature of *XPO1* was highly expressed and inversely related to left ventricular function in heart failure patients of ischemic etiology. Therefore, we hypothesized that in a rat model of myocardial infarction treated with AAV9-shXPO1, an improvement in the ventricular function after a follow-up period may be observed.

**Methods:** We induced myocardial infarction by coronary ligation in Sprague-Dawley rats (n=10), five of them received AAV9-shXPO1 treatment after four months and the other five infarcted rats did not receive any treatment. For non-failing controls, healthy Sham rats (n=5) received the placebo AAV9-scramble. Serial echocardiographic assessment was performed before, as well as, two and five months after intravenous injection.

**Results:** AAV9-shXPO1-treated rats showed improved fractional shortening (16.8 ± 2.8 *vs* 24.6 ± 4.1%, *P*<0.05) and LV systolic (5.10 ± 0.79 *vs* 3.52 ± 0.88mm, *P*<0.05) and diastolic (6.17 ± 0.95 *vs* 4.70 ± 0.93mm, *P*<0.05) diameters when comparing measurements obtained before and five months after AAV9 injection. We did not observe this improvement in untreated infarcted rats. Furthermore, EXP-1 levels in rats heart, brain, skeletal muscle, and liver were determined by western-blot and compared to controls in AAV9-shXPO1-treated rats. Lower levels of EXP-1 in cardiac tissue were observed (*P*<0.05).

**Discussion:** At five months follow-up, ischemic AAV9-shXPO1-treated rats showed partial recovery of LV myocardial function. No secondary symptoms attributable to AAV9-shXPO1 were observed in skeletal muscle, liver and brain.

## Introduction

Heart failure (HF) is a multifactorial condition accompanied by changes in ventricular function and dilation of cardiac chambers, causing increased morbidity, mortality and healthcare costs. This condition is closely associated with molecular and cellular alterations [1-4], such as the existence of various alterations in the molecular machinery of nuclear-cytoplasmic transport [5], which precisely regulates the bidirectional selective protein flow between the nucleus and the cytoplasm.

Previous studies have shown that several molecules that participate in nuclear-cytoplasmic transport (Exportin-1 [EXP-1], IMP-β3, Nup160) are intimately related to a reduced left ventricular (LV) function in human HF. It has been found the transcriptomic signature of these alterations and has been identified that changes in gene expression, specifically of *XPO1* that encodes EXP-1, were highly related to LV dysfunction in patients with HF of ischemic etiology [5, 6].

The short hairpin RNA (shRNA) can be used to silence specific genes and is a powerful tool in studies pertaining to loss of gene function and characterization. The use of highly cardio-tropic adeno-associated virus vector (AAV) with high affinity for the heart and down to other organs, can be introduced simply by intravenous injection [7, 8]. In particular, AAV9 has a great potential as a valuable tool for cardiac therapy in HF experimental models for RNA interference [9] and gene therapy [10].

We hypothesize that manipulation of gene deregulation has therapeutic value in HF. We aim to investigate whether highly significant relationship between *XPO1* and ventricular function is a component of causality. Therefore, we have developed a rodent ischemic heart failure experimental model to show whether AAV9-shXPO1 silencing agent induces recovery of myocardial function.

## Materials and Methods

### Ethics statement

The project was approved by the Biomedical Investigation Ethics Committee of La Fe Hospital. The investigation conforms to the *Guide for the Care and Use of Laboratory Animals* published by the US National Institutes of Health (NIH Publication No. 85-23, revised 1996), and the National (RD 53/2013) and European Directive (2010/63/EC). All surgery was performed using accurate anesthesia and analgesia veterinary protocols, to minimize animal suffering. All animal procedures are reported following ARRIVE (Animal Research: Reporting of *In Vivo* Experiments) guidelines and included as Supplementary Data (S1).

### Rat heart failure model

Adult male Sprague-Dawley rats weighting 300-400 g were anesthetized and left anterior descending (LAD) coronary artery ligation was performed (n=10) with Polypropylene non-absorbable sutures (Premilene^®^ Braun) to induce chronic myocardial infarction. Four months later, a group of rats with reduction of heart function received intravenous AAV9-shXPO1 gene therapy (n=5), and another did not receive any treatment (n=5) (see below). Age-matched non-infarcted sham rats (n=5) served as healthy non-failing controls (CNT) and received intravenous placebo AAV9-scramble. The experimental design of these groups and the experimental procedure (Fig 1) was performed based on previous studies focused on this field of research [11-13].

**Fig 1.**
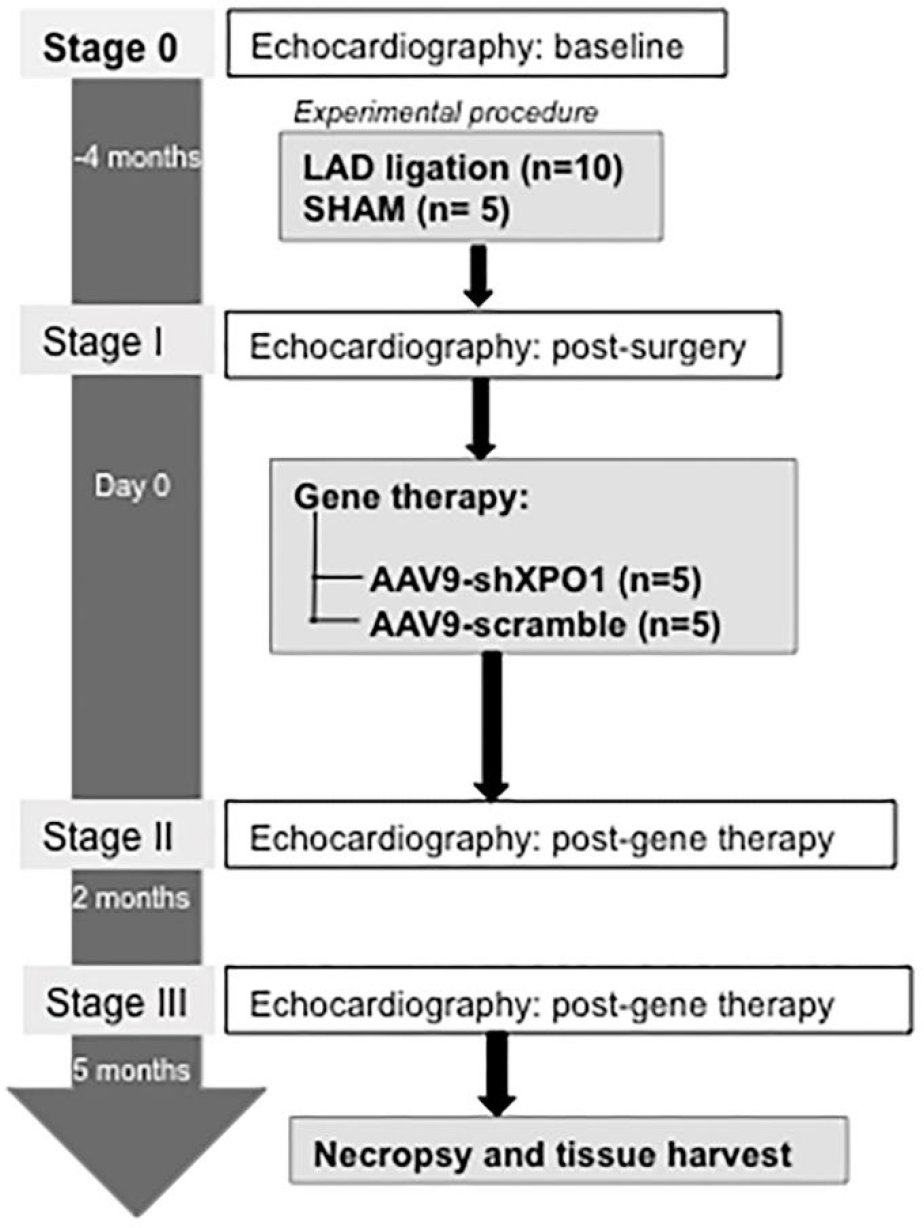
Study protocol timeline. Baseline functional echocardiographic parameters were evaluated (Stage 0) in rats before the experimental procedure. Functional parameters were studied in SHAM and LAD (left anterior descendent coronary artery) ligation animals after procedure (Stage I). Four months later, animals were administered with either Adeno-associated virus vector silencing *XPO1* gene (AAV9-shXPO1) or placebo (AAV9-scramble) particles. After gene therapy, changes in functional parameters were evaluated by echocardiography two (Stage II) and five (Stage III) months later.

### Gene therapy

AAV9-shXPO1 and placebo AAV9-scramble were manufactured by Creative Biogene (Shirley, NY). Based on previous literature, different AAV9 particles were administered (5 x 10^11^ genomes) [9], by tail vein injection at 16 weeks after infarction in infarcted rats (AAV9-shXPO1) and in non-infarcted rats (AAV9-scramble). Five months after vector delivery, the rats were sacrificed and their organs (heart, brain, skeletal muscle, and liver) were explanted. All samples were stored at - 80°C until protein extraction.

### Echocardiographic assessment

A noninvasive transthoracic echocardiographic method was used to evaluate the morphology and function of the left ventricle. Echocardiography was performed on anesthetized animals with ketamine and valium (26 and 6 mg/kg, respectively) and so that their anterior chest was shaved. Heart Functional parameters were analyzed with a two-dimensional mode using Philips EnVisor M2540A Ultrasound System before the LAD ligation and AAV9-shXPO1 administration as well as two and five months after the gene therapy.

### Tissue sampling

Frozen samples (50 mg of heart, lung, brain, skeletal muscle and liver) were homogenized in a total protein extraction buffer (2 % SDS, 10 mM EDTA, 6 mM Tris–HCl, pH 7.4) with protease inhibitors (25 μg/mL aprotinin and 10 μg/mL leupeptin) in a FastPrep-24 homogenizer with specifically designed Lysing Matrix D tubes (MP Biomedicals, USA). The homogenates were centrifuged and the supernatants were aliquoted. The protein content of the aliquots was determined by Peterson’s Modification [14] of the Lowry method using bovine serum albumin (BSA) as standard.

### Western blot analysis

Protein samples for detection of EXP-1 and GAPDH were separated using Bis-Tris Midi gel electrophoresis with 4–12% polyacrylamide under reducing conditions. Description of western blot procedure are extensively described by Gil-Cayuela et al [15]. The primary detection antibodies used were anti-Exportin-1 (611833) mouse monoclonal antibody (1:50) from BD Transduction Laboratories™, and anti-GAPDH (ab9484) mouse monoclonal antibody (1:1000) obtained from Abcam and used as a loading control.

The bands were visualized using an acid phosphatase-conjugated secondary antibody and nitro blue tetrazolium/5-bromo-4-chloro-3-indolyl phosphate (NBT/BCIP, Sigma-Aldrich, St. Louis, USA) substrate system. Finally, the bands were digitalized using an image analyzer (DNR Bio-Imagining Systems, Israel) and quantified with the GelQuant Pro (v. 12.2) program.

## Statistical methods

The Kolmogorov-Smirnov test was used to analyze the distribution of the variables. All variables were normally distributed. Data are presented as mean value ± standard deviation. Comparisons of variables were analyzed using Student’s t-test. Significance was accepted at the *P* < 0.05 level. All statistical analyses were performed using SPSS software v. 20, 2012 for Windows (IBM SPSS Inc., Chicago, IL, USA).

## Results

### *XPO1* silencing and AAV9-shXPO1 specificity

To determine the efficacy of *XPO1* silencing and specificity of the AAV9-shXPO1 vector, we measured EXP-1 levels in different explanted tissues of rats by western blot. Compared to the CNT group, the treated group showed lower EXP-1 levels in cardiac tissue (100 ± 16 vs 76 ± 9 arbitrary units, au, *P* < 0.05) (Fig 2A). Both the CNT and the treated group had similar EXP-1 levels in the skeletal muscle, liver, and brain (100 ± 31 vs 109 ± 26 au, 100 ± 28 *vs*103 ± 25 au and 100 ± 14 vs 110 ± 17 au, *P* > 0.05; respectively) (Fig 2B, C and D). Additionally, no secondary effects produced by the AAV9-shXPO1 vector were observed.

**Fig 2.**
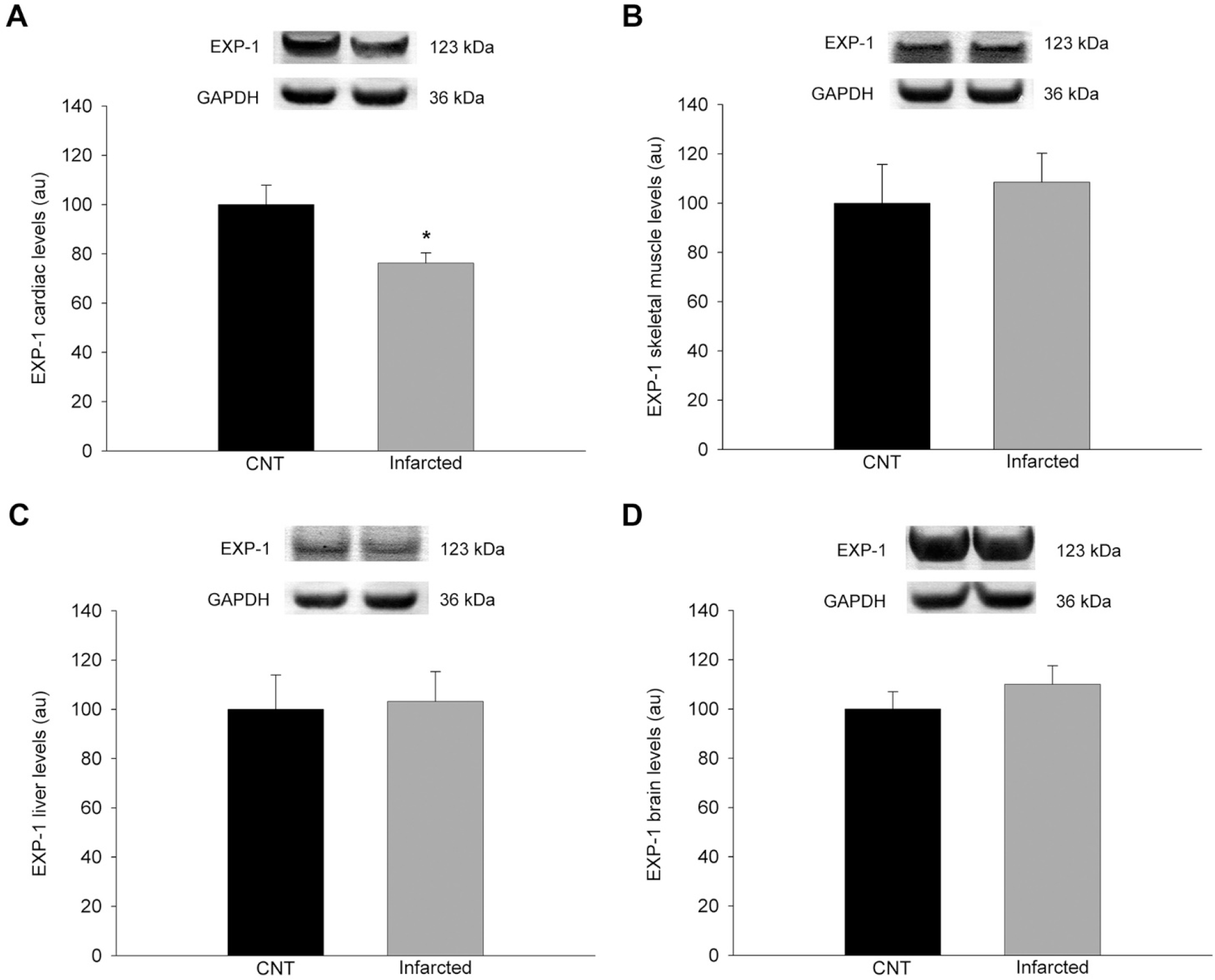
EXP-1 levels in infarcted and healthy non-failing control rats. (A) Heart. (B) Skeletal muscle. (C) Liver. (D) Brain. The values are normalized to GAPDH, and then to the CNT group. The values from the CNT group were set to 100. The data are expressed as mean ± SEM in arbitrary units. CNT, controls; au, arbitrary units. \**P* < 0.05.

### Echocardiographic assessment

Cardiac function in infarcted and CNT rats was measured by noninvasive transthoracic echocardiography to evaluate ventricular function and diameters. Echocardiographic measurements were taken prior to surgery and treatment of rats (stage 0). Fractional shortening (FS) of infarcted rats group (prior to surgery and injection) was 31.3 ± 8.6%, and LV end-systolic (LVESD) and LV end-diastolic (LVEDD) diameters were 2.57 ± 0.60 mm and 3.73 ± 0.63 mm, respectively. Infarcted rats without treatment group (prior to surgery) had a FS of 30.8 ± 7.6%. CNT rats had normal FS (31.1 ± 8.0%), and LVESD and LVEDD (2.53 ± 0.61 mm and 3.75 ± 0.59 mm, respectively).

After coronary ligation (stage I), FS of infarcted rats was 16.8 ± 2.8%, and LVESD and LVEDD were 5.10 ± 0.79 mm and 6.17 ± 0.95 mm, respectively. Infarcted rats without treatment had a FS of 16.7 ± 2.7%. CNT rats had normal FS (31.3 ± 8.5%).

At stage II (two months after AAV9 injection), treated infarcted rats had a FS of 16.4 ± 2.4%, LVESD of 5.03 ± 0.87 mm, and LVEDD of 6.14 ± 1.14 mm. Infarcted rats without treatment had a FS of 16.5 ± 2.5% and CNT rats remained with normal echocardiographic parameters respect to the beginning of the study. At stage III (five months after AAV9 injection) treated infarcted rats had a FS of 24.6 ± 4.1% (*P* < 0.05 compared to stage I, Fig 3A), LVESD of 3.52 ± 0.88 mm (*P* < 0.05 compared to stage I, Fig 3B), and LVEDD of 4.70 ± 0.93 mm (*P* < 0.05 compared to stage I, Fig 3C). Infarcted rats without treatment had a FS of 16.5 ± 2.6%. CNT rats maintained their parameters.

**Fig 3.**
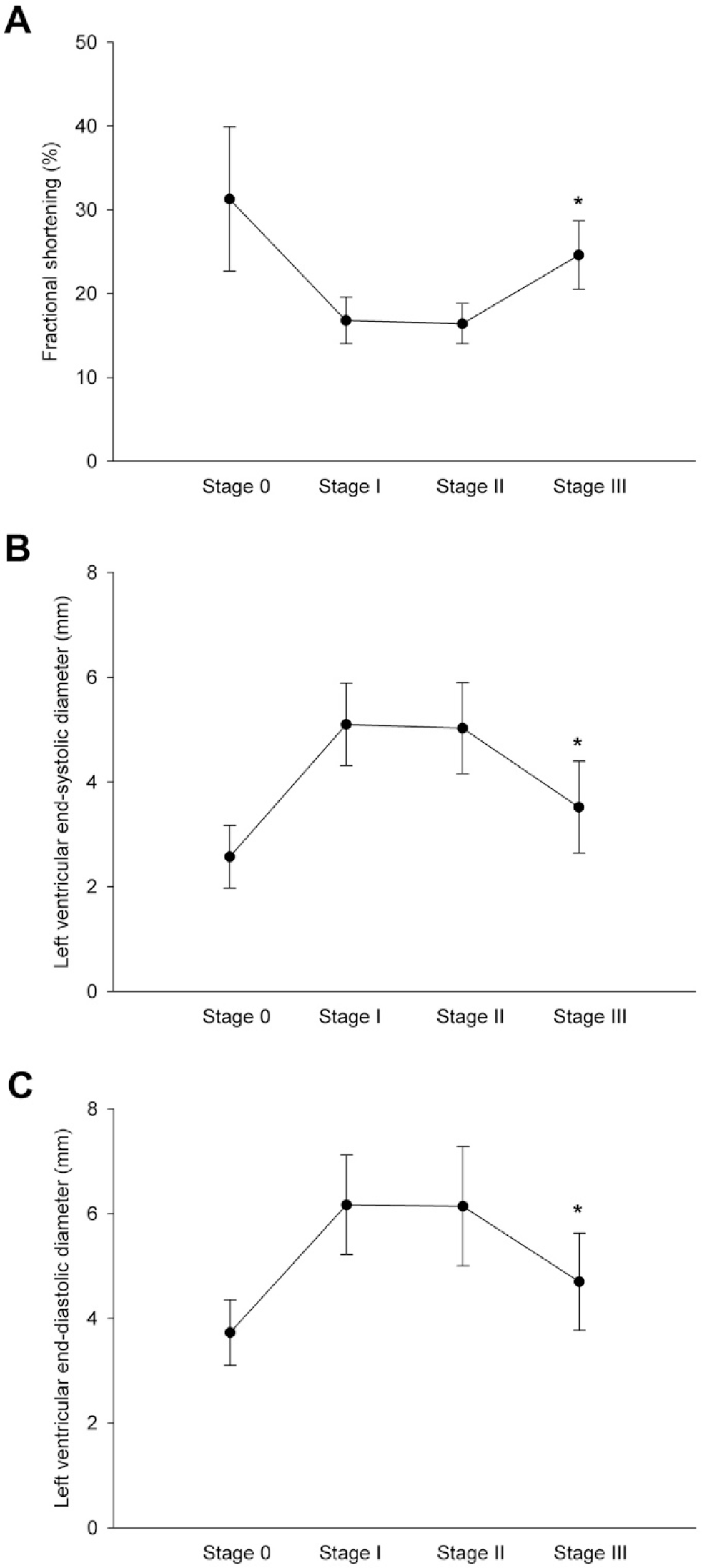
Echocardiographic parameters of infarcted treated rats measured before surgery and injection (Stage 0), before injection (Stage I), and two (Stage II) and five months (Stage III) after AAV9-shXPO1 injection. (A) Fractional shortening. (B) Left ventricular end-systolic diameter. (C) Left ventricular end-diastolic diameter. \**P* < 0.05 Stage I vs Stage III.

## Discussion

This study may provide a new therapeutic strategy based on gene therapy to restore ventricular function in patients with ischemic cardiomyopathy, which is the leading cause of death worldwide and lacking effective treatment [16]. In this study, we report a successfully delivery of an AAV-based gene therapy in a long-term chronic HF rat model, which simulates the clinical features seen in patients with chronic HF from an ischemic origin. We have expanded the follow-up until 5 months after gene delivery, proving the long-term efficacy of the treatment and showing the safety of this procedure, since there was no evidence of side-effects in the animals.

EXP-1 mediates the nuclear export of proteins, rRNA, snRNA, and some mRNAs. Previous studies in patients with ischemic cardiomyopathy showed elevated expression levels of both EXP-1 mRNA and protein, and interestingly, these levels were inversely related with ejection fraction and positively correlated with LVESD and LVEDD [5, 6], *i.e*. higher EXP-1 expression is linked with LV function impairment. Hence, we intended to demonstrate the cause-effect of this relationship in this study through gene therapy, which may result in a useful approach for the treatment of heart disease [10, 17, 18]. Cardiac gene therapy uses vectors that can robustly, specifically, and persistently deliver therapeutic genetic materials to the heart without generating local and/or systemic toxicity. The adenoviral vectors represent an efficient but unstable gene delivery vector for the heart. Nonetheless, long-term myocardial transduction in adult animals has been accomplished with the development of AAV vectors [19]. Research has established AAV9 as a cardio-tropic vector superior to all the other serotypes in rodents, making it the most appropriate vector for gene delivery to the heart [7, 8]. Our results support this property of AAV9, since we observed that the cardiac tissue was the only sample analyzed where EXP-1 levels decreased in infarcted AAV9-shXPO1 rats compared to that in the CNT rats. Furthermore we show the effectiveness and stability of the vector AAV9-shXPO1 as EXP-1 levels decreased in heart tissue five months after transduction.

Left ventricular function parameters are directly related to ventricular remodeling that occurs after injury of the heart muscle. Ventricular function of infarcted rats appeared to be in partial recovery following *XPO1* silencing. At two months after injection (Stage II) cardiac function in the rats was similar to that immediately after coronary ligation (Stage I). Nevertheless, at five months after injection, the differences in these parameters were significant and resulted in improvements in the state of cardiac function in infarcted rats with *XPO1* silencing. This silencing could have similar effects at revascularization, recovering the hibernating myocardium and thereby improving ventricular function in infarcted rats. Although, this partial recovery would be slower than that achieved by performing a bypass procedure, it is less invasive and harmful. We did not observe an improvement in the cardiac function of untreated infarcted rats; these rats maintained similar parameters during the follow-up after coronary ligation.

## Study limitations

In order to rationalize funding resources and minimize animal testing, we decided to compare only gene therapy responses in rats with chronic infarction, not studying AAV9-shXPO1 administration in healthy control rats [11]. For the development of the heart failure model, a standardized protocol of veterinary pharmacology and surgery was followed, still, the manual procedure of coronary ligation may introduce some variability between infarcted rats.

## Conclusions

In conclusion, after the five-month follow-up, infarcted rats treated with AAV9-shXPO1 displayed partial recovery of LV myocardial function. No secondary AAV9-shXPO1-attributable symptoms were observed in the cardiac-related organs. In this study we have developed a heart failure rat model that have been treated with AAV9-shXPO1, producing the gene silencing of XPO1. After a follow-up period, we have observed an improvement in ventricular function of these rats. This study provides a new therapeutic strategy based on gene therapy to restore ventricular function in patients with ischemic cardiomyopathy.

## Abbreviations

AAV: adeno-associated virus vector
EXP-1: Exportin-1
FS: Fractional shortening
HF: Heart failure
LAD: left anterior descending
LV: left ventricular
LVEDD: left ventricular end-diastolic diameter
LVESD: left ventricular end-systolic diameter
shRNA: short hairpin RNA

## Acknowledgment

The authors thank the Animal Experimentation Platform of the IIS La Fe, for the technical support in this work. This work was supported by the National Institute of Health *Fondo de Investigaciones Sanitarias del Instituto de Salud Carlos III* [PI16/01627, PI17/01925, PI17/01232], *Consorcio Centro de Investigación Biomédica en Red, M.P.* [CIBERCV, under Grant CB16/11/00261], and the European Regional Development Fund (FEDER).

## Supplementary Data (S1)

### Protocol. Experimental procedure in rats

#### Myocardial infarction

##### Anesthesia and analgesia

To perform the infarction in the animals, a standardized anesthetic protocol was used. The first step consisted in the induction of the animal kept in an anesthetic chamber with a mixture of oxygen and sevoflurane (5% v/v Abbott) for approximately 2 minutes or until a cessation of motor activity was observed. Endotracheal intubation was carried out with an 18 G (1.3 mm diameter) catheter and a 2 “(5.1 cm) length (Braun). For this, the rat was placed in the supine position on a platform and the limbs were clamped to stabilize the position. By lighting with a cold light source directed to the ventral area of the neck and gently extracting the tongue, the glottis and trachea of the rat were visualized. Intubation was performed and the rat was immediately connected to an automatic respirator (Harvard Apparatus^®^, model 683) with set parameters of 2 ml of tidal volume and 140 breaths per minute. The correct intubation was checked by confirming breath on a glass surface first and ensuring rhythmic movement of the chest with respect to assisted ventilation afterwards. The tube was fixed to the animal and to the surgical table. The appropriate anesthetic level was maintained during the surgical procedure with 2% sevofluorane, 0.3 ml of N_2_O and 0.1 ml of O_2_. The rat was placed on a cork surface and covered with a surgical drape. An analgesic drug, fentanyl (Fentanest^®^, Kern Pharma) was used at a dose of 0.05 mg/kg inoculated immediately before starting surgery intraperitoneally (IP) as well as a dose of 0,05 mg/kg IP of buprenorphine (Buprex^®^, Schering-Plow SA) 20 minutes before procedure. In order to avoid muscle contractions, 6 mg/kg lidocaine dose was infiltrated intraincisionally at the beginning of the procedure. In the post-surgical protocol, animals were given a dose of 0.05 mg/kg of buprenorphine IP every 6-8 hours for 48 hours. The euthanasia of the rats was carried out according to the regulations in force, administering an anesthetic dose of ketamine/diazepam and performing a subsequent exsanguination, or inoculating 1 ml of intracardiac 0.2 M KCl.

##### Surgical procedure

The rat was placed in the supine position on an isolated cork base and the limbs were immobilized with adhesive tapes. The eyes were hydrated with a gel tear to avoid corneal ulcers (Lubrifilm^®^, Alcon Cusí). For the preparation of the surgical area, the thorax was shaved and disinfected with diluted povidone (Betadine^®^, Viatris Manufacturing). Animals were covered with a surgical drape leaving the thorax free to perform a lateral thoracotomy between the 4th and 5th left intercostal space.

First, a two-centimeter skin incision was made directed towards the sternum and middle area of the thorax. The subcutaneous tissue was separated and after the dissection of the pectoral and intercostal muscles (the latter cranially to the rib to avoid the vascular bundle that runs caudally to them) was accessed to the thoracic cavity. By placing spacers in the intercostal space and gauze below the thorax of the rat it was possible to expose the heart. Then, with the help of a swab soaked in physiological saline, the thymus was removed cranially. Then, with gentleness and avoiding the cranial lung lobe, pericardiotomy was performed incising with blunt microsurgical forceps. The area was cleaned with a wet and sterile gauze to be able to visualize the left anterior descending coronary artery and ligation area was delimitated starting from about 4 mm below the left atrium. For the occlusion, 5/0 non-absorbable monofilament suture (Premilene^®^, Braun) was used by permanent ligation. Correct ligation of the artery and infarction was evidenced, due to the change that occurs immediately in the infarcted area (bluish color).

Finally, the thorax was sutured with a 3/0 resorbable monofilament suture (Monosyn^®^, Braun), ensuring that there was no thoracic permeability so that negative chest pressure recovered after closure. The muscular layer was closed with a continuous suture and the tissues were irrigated with saline. Finally, the skin was closed with an inthermal suture and the wound was disinfected with povidone. A few minutes after removing the inhalation agent they began to see respiratory reflexes and ventilations of the animal, at which point it was disconnected and left isolated and tempered in a cage with an electric blanket.

##### Echocardiographic asessment

To evaluate the ventricular function of the rats, previous and postoperative echocardiograms were performed. In this way, detailed information on the cardiac morphology of each individual and its evolution over time was obtained. A combination of anesthetic drugs was administered for the immobilization of the animals and to be able to take the images. A dose of 26 mg/kg of ketamine (Ketalar^®^, Pfizer) and 6 mg/kg of diazepam (Valium^®^, Roche) was prepared and 15 minutes prior to echocardiography half of the dose was inoculated to the rat, inoculating the remaining one if was necessary. The thorax was shaved and the animal was placed on an electric blanket in supine position. For the acquisition of images, the Philips EnVisor M2540A Ultrasound System was used together with a linear 10 MHz transducer. Imaging was performed 1 to 3 days before surgery, to record the baseline ventricular function of the rats, post-infarction was performed 1 week, and 2 and 5 months after gene therapy. In the study, transthoracic echocardiography was performed on the short axis with 3 cuts: one behind the atria, another cut in the middle ventricular area at the level of the papillary muscles and finally another in the apex area. Images were taken in M-mode (linear image to which the time variable has been introduced) and also in 2D mode (two-dimensional image in real time). To calculate the functional parameters, three consecutive cardiac cycles were measured using standard methods. The diameters of the left ventricle were measured in systole and diastole in the M-mode and the internal area in the 2D mode. From these measurements, the shortening fraction and the ejection fraction were calculated.

